# *Salmonella enterica* serovar Typhimurium ST313 responsible for gastroenteritis in the UK are genetically distinct from isolates causing bloodstream infections in Africa

**DOI:** 10.1101/139576

**Authors:** Philip M. Ashton, Sian V. Owen, Lukeki Kaindama, Will P. M. Rowe, Chris Lane, Lesley Larkin, Satheesh Nair, Claire Jenkins, Elizabeth de Pinna, Nicholas Feasey, Jay C. D. Hinton, Tim Dallman

## Abstract

The ST313 sequence type of *Salmonella enterica* serovar Typhimurium causes invasive non-typhoidal salmonellosis amongst immunocompromised people in sub-Saharan Africa (sSA). Previously, two distinct phylogenetic lineages of ST313 have been described which have rarely been found outside sSA. Following the introduction of routine whole genome sequencing of *Salmonella enterica* by Public Health England in 2014, we have discovered that 2.7% (79/2888) of S. Typhimurium from patients in England and Wales are ST313. Of these isolates, 59/72 originated from stool and 13/72 were from extra-intestinal sites. The isolation of ST313 from extra-intestinal sites was significantly associated with travel to Africa (OR 12 [95% CI: 3,53]). Phylogenetic analysis revealed previously unsampled diversity of ST313, and distinguished UK-linked isolates causing gastroenteritis from African-associated isolates causing invasive disease. Bayesian evolutionary investigation suggested that the two African lineages diverged from their most recent common ancestors independently, circa 1796 and 1903. The majority of genome degradation of African ST313 lineage 2 is conserved in the UK ST313 lineages and only 10/44 pseudogenes were lineage 2-specific. The African lineages carried a characteristic prophage and antibiotic resistance gene repertoire, suggesting a strong selection pressure for these horizontally-acquired genetic elements in the sSA setting. We identified an ST313 isolate associated with travel to Kenya that carried a chromosomally-located *bla*_CTX-M-15_, demonstrating the continual evolution of this sequence type in Africa in response to selection pressure exerted by antibiotic usage.

The S. Typhimurium ST313 sequence type has been primarily associated with invasive disease in Africa. Here, we highlight the power of routine whole-genome-sequencing by public health agencies to make epidemiologically-significant deductions that would be missed by conventional microbiological methods. The discovery of ST313 isolates responsible for gastroenteritis in the UK reveals new diversity in this important sequence type. We speculate that the niche specialization of sub-Saharan African ST313 lineages is driven in part by the acquisition of accessory genome elements.

## Introduction

Serovars of *Salmonella enterica* cause infections in a diverse range of hosts. In humans, Salmonellae are responsible for a broad range of clinical presentations, from gastroenteritis to invasion of normally sterile compartments such as the bloodstream or brain. Two serovars, *Salmonella* Typhi and *Salmonella* Paratyphi A are particularly associated with both human-restricted, and invasive disease. The clinical syndrome caused by these serovars is known as Typhoid or Enteric fever, and this has led to the remaining 2,600 serovars being loosely described as non-typhoidal. By inference, “non-typhoidal” serovars have been considered to be non-invasive in immunocompetent individuals; this crude clinical distinction is misleading (1).

These “non-typhoidal” *Salmonella* (NTS) serovars typically have a broad host-range, and the majority of human cases are foodborne, often originating from zoonotic reservoirs (2). Whilst most infections are typically associated with self-limiting gastroenteritis, in a minority (∼5%) invasive disease occurs, frequently due to human host immunosuppression, for example advanced HIV infection (3). NTS are a significant public health burden worldwide and *S*. Typhimurium and *S*. Enteritidis are the predominant serotypes observed in clinical cases in most countries. In England & Wales, 48.7% of the isolates referred to the *Salmonella* Reference Service were *S*. Typhimurium or *S*. Enteritidis (PHE figures) in 2012.

The clinical distinction between typhoidal and nontyphoidal disease is particularly unhelpful in sub-Saharan Africa (sSA), where non-typhoidal serovars are amongst the most common cause of bloodstream infection, a clinical condition known as invasive nontyphoidal *Salmonella* (iNTS) disease (1). Whilst the high prevalence of immunosuppressive illness such as HIV and malaria in sSA are clear predisposing factors for the emergence of iNTS disease as a major public health problem, the huge burden of disease has led to further investigation into the serovars responsible for iNTS disease. *S*. Typhimurium is the serovar most commonly associated with this condition (4).

Multi locus sequence typing (MLST) is a molecular approach for typing micro-organisms, and uses the allelic varation of seven highly conserved *Salmonella* housekeeping genes to approximate bacterial phylogeny (5). Whole genome sequence studies of isolates collected from patients with iNTS disease in sSA initially identified a novel sequence type (ST), ST313, of *S.* Typhimurium in 2009 (6). Subsequent genome-based studies confirmed two distinct phylogenetic lineages of ST313, and spatio-temporal phylogenetic reconstruction suggested that lineage 1 emerged around 1960 in south west Africa, whereas lineage 2 emerged around 1977 in Malawi (5). Both lineages are associated with antimicrobial resistance (AMR) mediated by differing Tn21-like integrons on the virulence plasmid pSLT (6), and it was proposed that clonal replacement of lineage 1 by lineage 2 had occurred, driven by the acquisition of chloramphenicol resistance in lineage 2 (7).

It has been suggested that the link between *S*. Typhimurium ST313 and iNTS disease in sSA is that, compared with the generalist S. Typhimurium ST19, ST313 has adapted to an extra-intestinal/invasive lifestyle via genome degradation (6, 8). This would be consistent with the finding of an accumulation of pseudogenes in pathways associated with gastrointestinal colonization, as observed in host-restricted *Salmonella* serovars such as *S.* Typhi, and in *Yersinia pestis, Shigella* spp, *Mycobacterium leprae* and *Bordetella pertussis* (9–14). Another observation from comparative genomic studies, which supports the hypothesized enhanced virulence of ST313 includes the detection of novel prophages BTP1 and BTP5 (15) including the reported BTP1 phage-encoded putative virulence gene, *st313-td* (16). A number of phenotypic characteristics which distinguish ST313 from gastroenteritis-associated ST19 strains have also been described including a reduction in motility, flagellin expression, stationary-phase catalase activity and biofilm formation (17–19). Despite these phenotypes, it remains to be proven whether ST313 are intrinsically more invasive than ST19.

Since April 1st 2014, every presumptive *Salmonella* isolate received by the *Salmonella* Reference Service (SRS) of Public Health England (PHE) has been whole genome-sequenced (WGS) to allow identification, characterization and typing in one laboratory process (5, 20).

In this study we investigated the prevalence of ST313 in cases of laboratory-confirmed *S*. Typhimurium infections reported in England and Wales, obtained clinical data regarding the origin of isolates (faeces or blood) and contacted patients to determine whether infection was associated with travel to high-incidence areas such as sSA. We used a phylogenetic approach to place the UK-isolated ST313 into the evolutionary context of the African ST313 lineages. We then investigated the accessory genome, multi-drug resistance (MDR) determinants and the presence of pseudogenes in the UK isolates to shed light on the population structure and evolution of ST313 *S.* Typhimurium. We also compared key phenotypes of the UK-isolated ST313 with the African ST313 lineage.

## Results

### Epidemiology of ST313 in the UK

Between January 2014 and May 2016, 2,888 *S*. Typhimurium isolates were whole-genome sequenced by Public Health England, of which 79 (2.7%) were of multi-locus sequence type ST313. Whole genome sequence data were available for a further 363 *S*. Typhimurium isolates from 2012, of which 7 (1.9%) were ST313. Of these 86 ST313 isolates (Supplementary Table S1), 75 were derived from human patients (5 patients had two isolates sequenced, and one patient had 5 isolates sequenced), 1 from a dog and 1 was isolated from an unspecified raw food sample. The sample type was recorded for 72 of the 75 human patient isolates; 13 patients had one or more isolate from extra-intestinal sites (blood, pus or bronchial alveolar lavage) indicating iNTS disease, and 59 isolates were from faeces alone (indicating gastrointestinal infection). Travel information was available for 51 of the 75 human patients of whom 8 reported travel to sSA during the estimated disease incubation period and one adult male reported consuming food of West African origin in London.

Of the 51 patients with travel information, 48 had sample type recorded. Of the 8 patients who reported travel to Africa, 6 had extra-intestinal infections. In contrast, just 2 of 40 patients for whom travel information was available and did not report travel to Africa had extra-intestinal infection, showing that travel to Africa is significantly associated with iNTS disease in the UK (OR 57.0 [95% CI: 6.7, 484.8], p-value = 0.0002) (Table 1A).

**Table 1.**
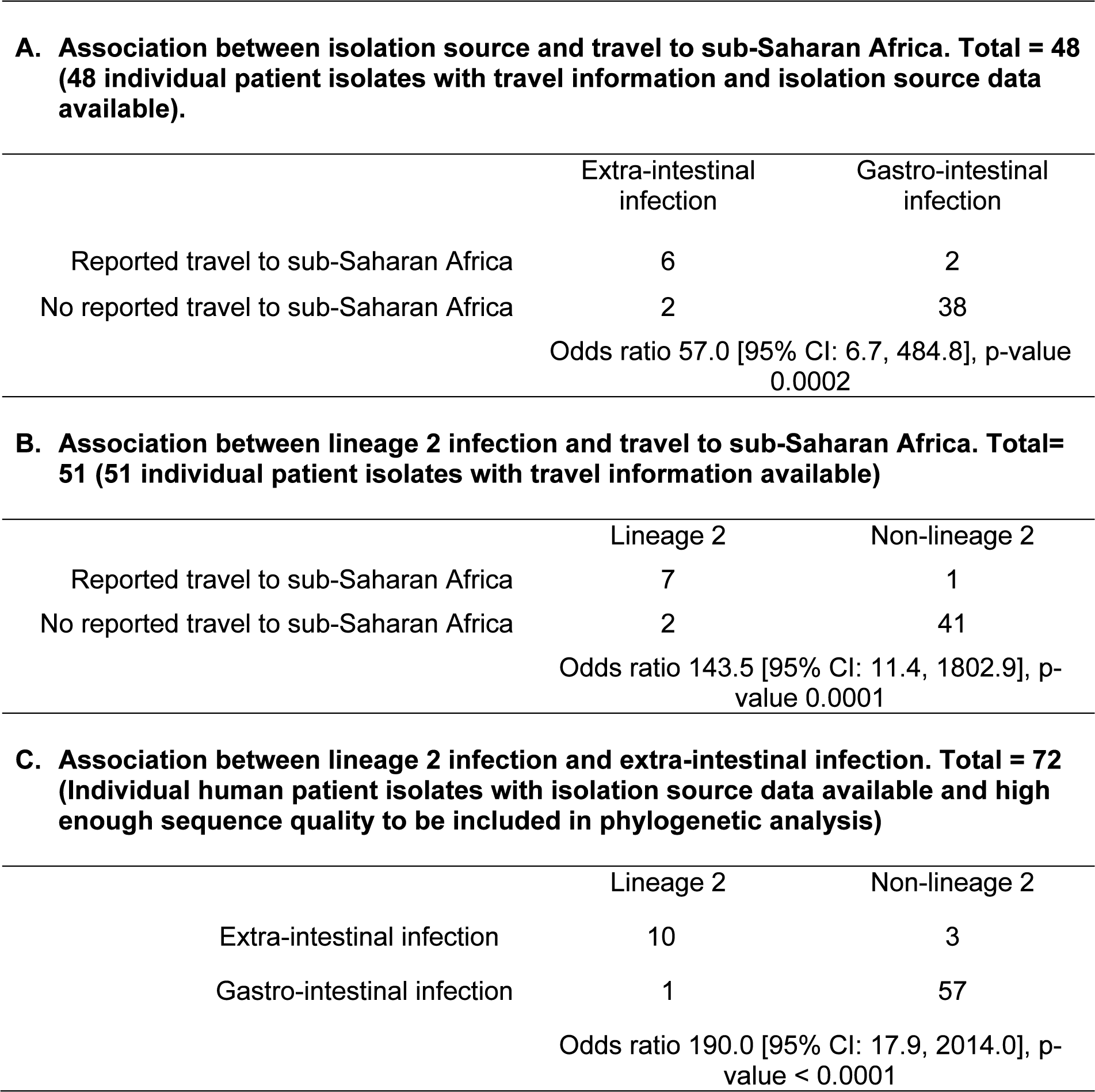
Summary of key epidemiological features of ST313 sampled by PHE. Full metadata for all isolates in this study is available in Supplementary Table S1.

### Phylogenetic analysis

Sequence data quality was sufficient to permit whole genome single nucleotide polymorphism (SNP) phylogenetic analysis of the isolates from 76 of the 77 patient and non-human isolates. Within the wider phylogenetic context of *S*. Typhimurium, all ST313 isolates submitted to PHE formed a monophyletic group that clustered with previously described African ST313 isolates (7) (Figure 1). A second maximum likelihood ST313 phylogeny was generated with a closely-related ST19 strain also received by PHE (strain U21) as an outlier, to study phylogenetic relationships (Figure 2A). Of the 76 isolates from distinct patients/sources 12 belonged to the previously described lineage 2, and 64 did not fall within any ST313 lineage that has been reported to date. No lineage 1 isolates were identified. Furthermore, the African associated lineages 1 and 2 did not form a monophyletic group within the novel diversity observed. Both of the African lineages share more recent common ancestors with UK-associated strains than with each other. Neither the food nor the animal isolate belonged to lineage 2. All the UK-derived isolates in this study originated from diagnostic laboratories in England and Wales. To simplify the categorization/differentiation of the lineages for discussion purposes, isolates belonging to lineage 1 and 2 (including those isolated in the UK) will be referred to as African lineages, and the non-lineage 1 and 2 isolates will be referred to as UK ST313. The UK ST313 isolates do not themselves form a coherent monophyletic cluster, revealing an unappreciated level of genetic diversity within ST313 (Figure 1).

**Figure 1.**
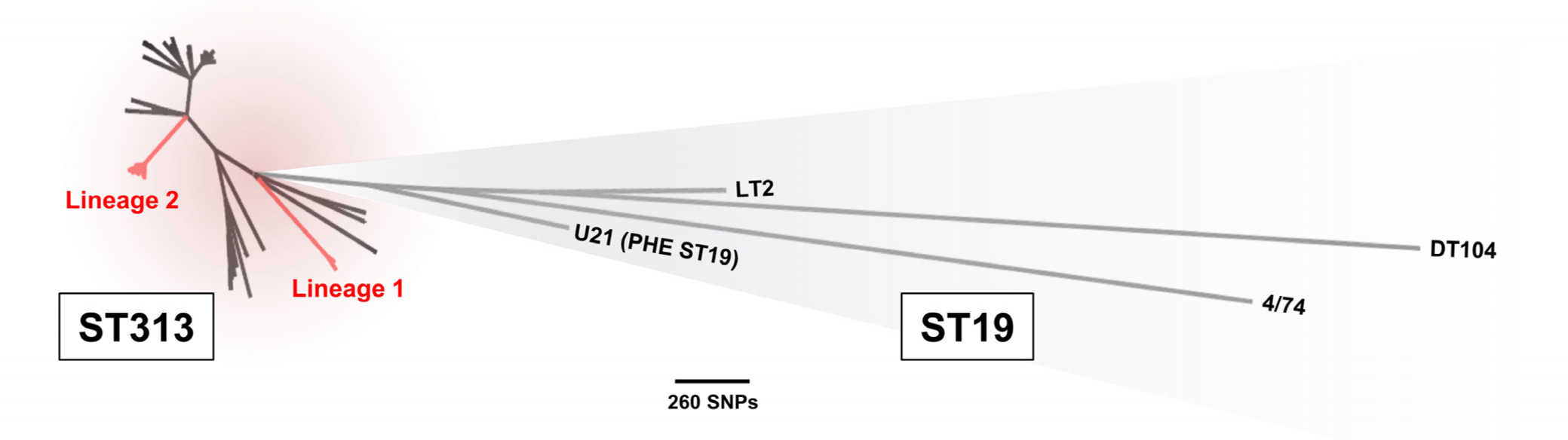
UK-ST313 isolates are phylogenetically distinct from African lineages. Unrooted maximum likelihood phylogeny of ST313 in the context of reference *Salmonella* Typhimurium ST19 isolates. Isolate U21 was an ST19 isolate, closely related to ST313, that was used as an outgroup in further ST313 analyses. The African epidemic ST313 lineages 1 and 2 are labelled.

To examine the evolutionary history of ST313, a maximum clade credibility tree was inferred using BEAST (Figure 3), and the topology was largely congruent with respect to the Maximum Likelihood tree (Figure 1, Figure 2). The most recent common ancestor (MRCA) of ST313 is estimated to have been in approximately 1787 (95% highest posterior distribution (HPD), 1735 - 1836). Lineage 1 diverged from other ST313 sampled in this study in approximately 1796 (95% HPD 1744-1842), while lineage 2 diverged from other ST313 sampled here in 1903 (95% HPD 1876 - 1927). The lineage 1 MRCA dated to around 1984 (95% HPD 1979-1987), while the lineage 2 MRCA dated to around 1991 (95% HPD 1986-1995). These confidence intervals overlap with the confidence intervals reported for the emergence of the two lineages by Okoro *et al*., 2012 (7). The two African lineages do not form a monophyletic group, and share an MRCA which is very close to that of ST313 as a whole. Both lineage 1 and 2 are separated from other sampled ST313 by long branches, indicating a distant MRCA with other isolates and suggesting that a tight population bottleneck has occurred relatively recently in evolutionary history.

**Figure 2.**
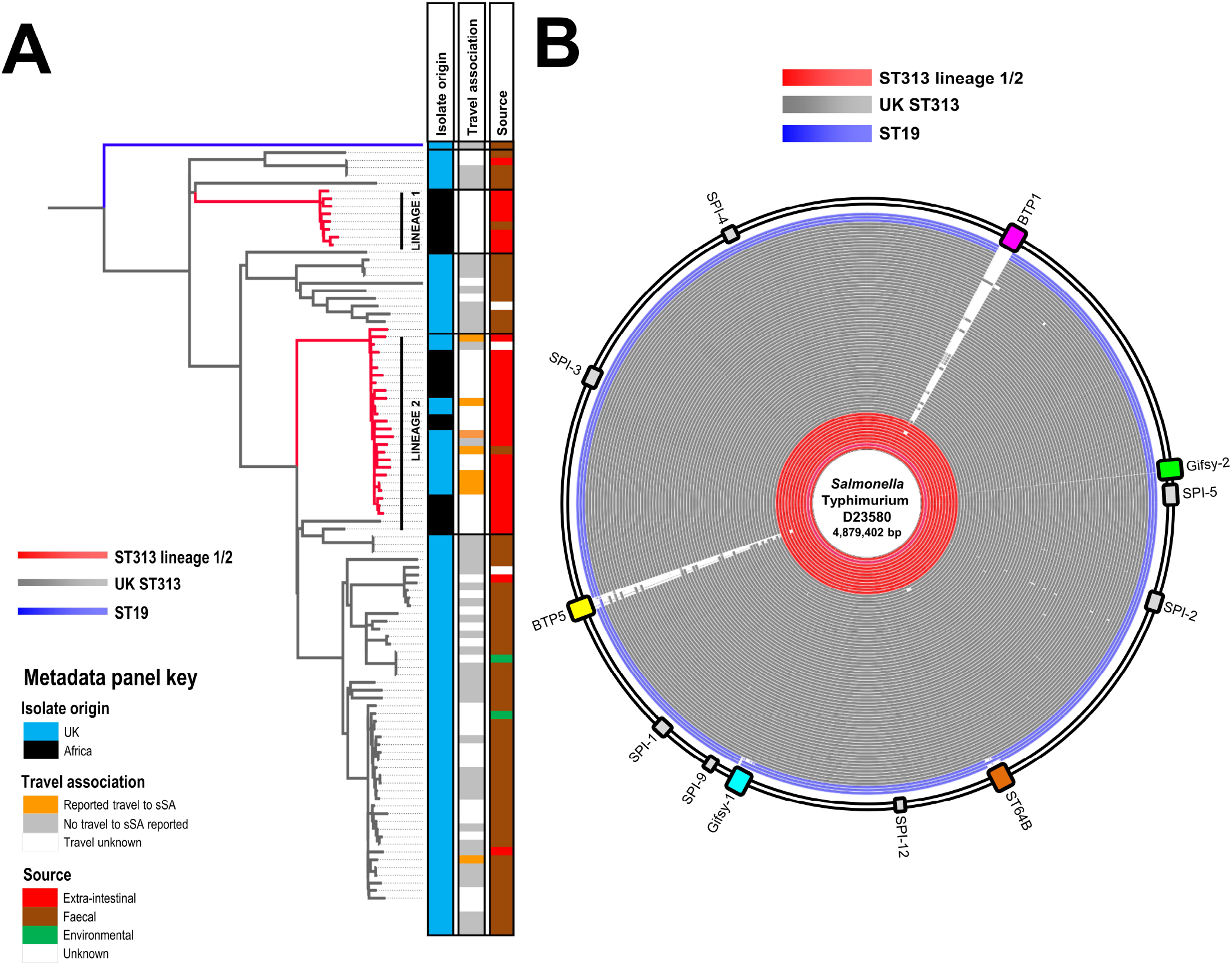
UK-ST313 are associated with gastrointestinal infection and do not harbour the African lineage associated prophages, BTP1 & BTP5. A. Maximum likelihood phylogeny of 77 UK-isolated ST313 strains received by PHE in the context of 24 African ST313 sequenced by Okoro *et al.* (2012). Red branches indicate ST313 lineage 1 and 2. Adjacent metadata panel showing: 1. Country isolates was associated with, Africa- orange, not Africa- blue; 2. Source, extra-intestinal- red, faecal- brown, environmental- green, unknown- grey. B. BLAST ring image showing BLAST comparison of all UK-isolated ST313 genomes (red and grey rings) along with 3 reference ST19 strains (blue rings) against lineage II representative strain D23580. The position of the prophages (coloured blocks) and *Salmonella* pathogenicity islands (grey blocks) in lineage II strain D23580 are shown around the outside of the ring.

**Figure 3.**
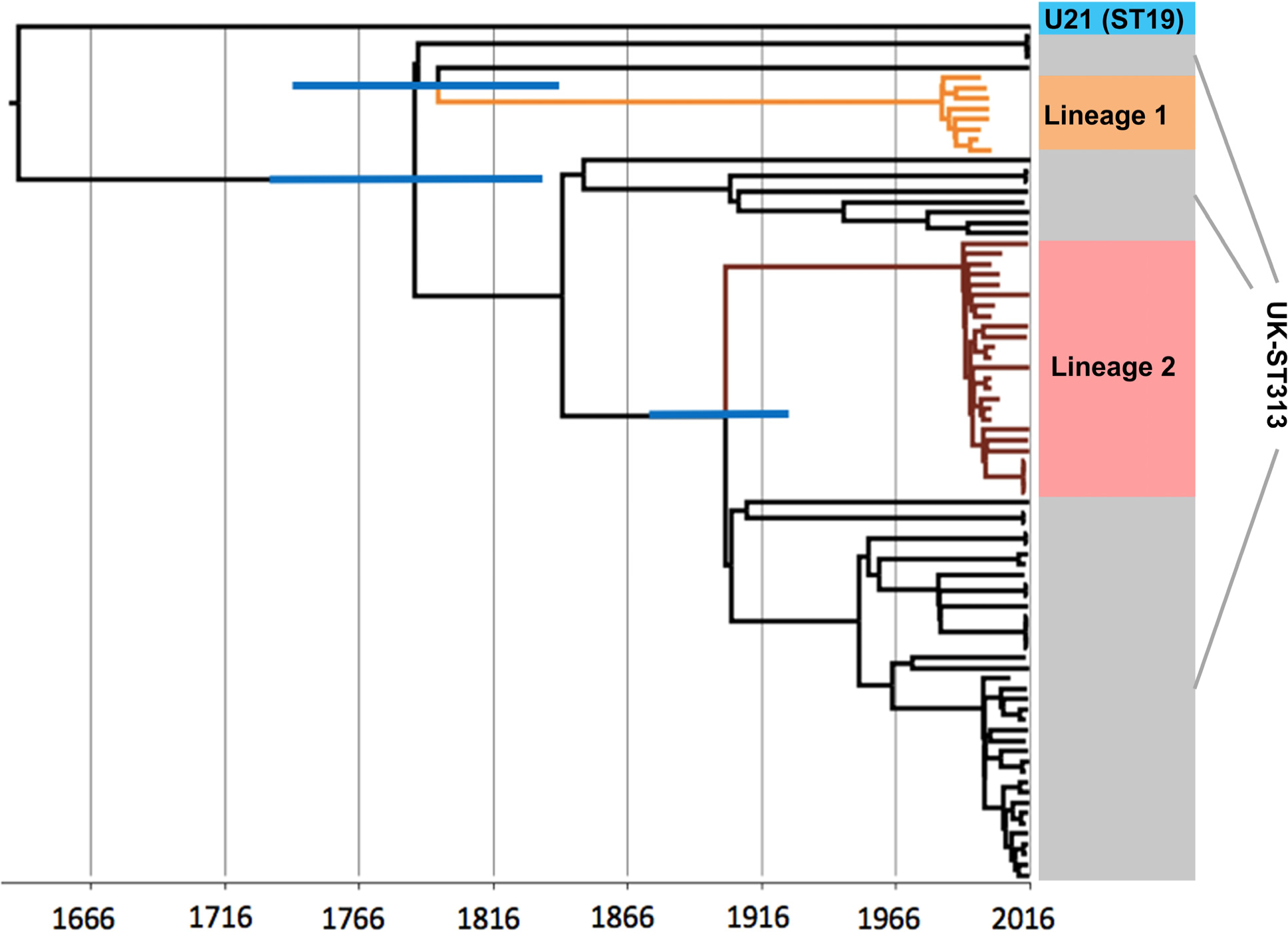
The timed phylogeny of all UK-isolated ST313 strains from this study and a representative sub-sample of ST313 genomes from Okoro *et al.*, 2012. Figure shows the maximum clade credibility tree from BEAST. Branches 95% HPD are displayed in blue for key nodes defining lineage 1 and lineage 2 (for tree with all 95% HPD, see supplementary Figure 4). Branches belonging to lineage 1 are coloured orange and branches belonging to lineage 2 are coloured brown.

### Epidemiological analysis in phylogenetic context

We investigated the association between reported travel to sSA and infection with the lineage 2 isolates. Of the 8 UK patients reporting travel to sSA during the seven days before disease onset, 7 were infected with a lineage 2 isolate. In contrast, of the 43 patients who reported no travel to sSA, 2 were infected with lineage 2 (Table 1B). This shows that travel to sSA was significantly associated with infection by ST313 lineage 2 (OR 143.5 [95% CI: 11.4,1802], p-value = 0.0001).

We investigated whether the ST313 lineage 2 isolates were more frequently associated with extra-intestinal or gastro-intestinal infection. Of the 11 patients infected with lineage 2 (and for which source of isolation data was available), 10 patients had isolates that originated from extra-intestinal sites. In contrast, for the patients infected with UK ST313 isolates, 3 of 60 isolates were from extra-intestinal sites (Table 1C). These data show that infection with lineage 2 is significantly associated with invasive disease (OR 190.0 [95% CI: 17.9, 2014.0], p-value < 0.0001).

### Accessory genome of ST313

Multi-drug resistance is a key phenotypic feature associated with both the African ST313 lineages, and is encoded by tn21-like integron insertions on the pSLT virulence plasmid. Analysis of the genome sequences indicated that all 76 ST313 isolates in this study carried the pSLT plasmid. However, the majority of the UK ST313 isolates were antibiotic-sensitive (59/64 were sensitive to all antimicrobials tested), and no consistent AMR gene profile was identified. In contrast, 10 of 12 UK-isolated lineage 2 isolates contained the same pSLT-associated MDR locus as the African lineage 2 reference strain D23580 (6). Four UK-isolated lineage 2 strains exhibited an atypical AMR gene profile; one isolate (U45) lacked the chloramphenicol resistance *catA* gene and another (U73) carried only the beta-lactamase gene *bla*_TEM1_. A third isolate, U1, had likely acquired resistance to fluoroquinolones via a mutation in the DNA gyrase subunit A gene, *gyrA*.

The last atypical UK-isolated lineage 2 isolate, U60, carried additional antibiotic resistance genes including extended-spectrum beta-lactamases (ESBL) *bla*_CTX-M-15_ and *bla*_OXA-1_, and genes conferring resistance to aminoglycosides, trimethoprim and tetracycline; *aac(6′)-Ib-cr*, *dfrA-14*, *tet(A)-1* (Figure 4a). Isolate U60 also encoded the tellurium heavy metal resistance operon (*terBCDEF*). Comparison to lineage 2 reference strain D23580 identified a putative 29kb deletion in the pSLT-BT virulence plasmid (6), which corresponded to the conjugal transfer region. Additionally, we identified sequence reads that mapped to 97% of the IncHI2 pKST313 plasmid, a novel plasmid which has recently been reported in lineage 2 isolates from Kenya and is known to encode ESBL resistance loci (21).

**Figure 4.**
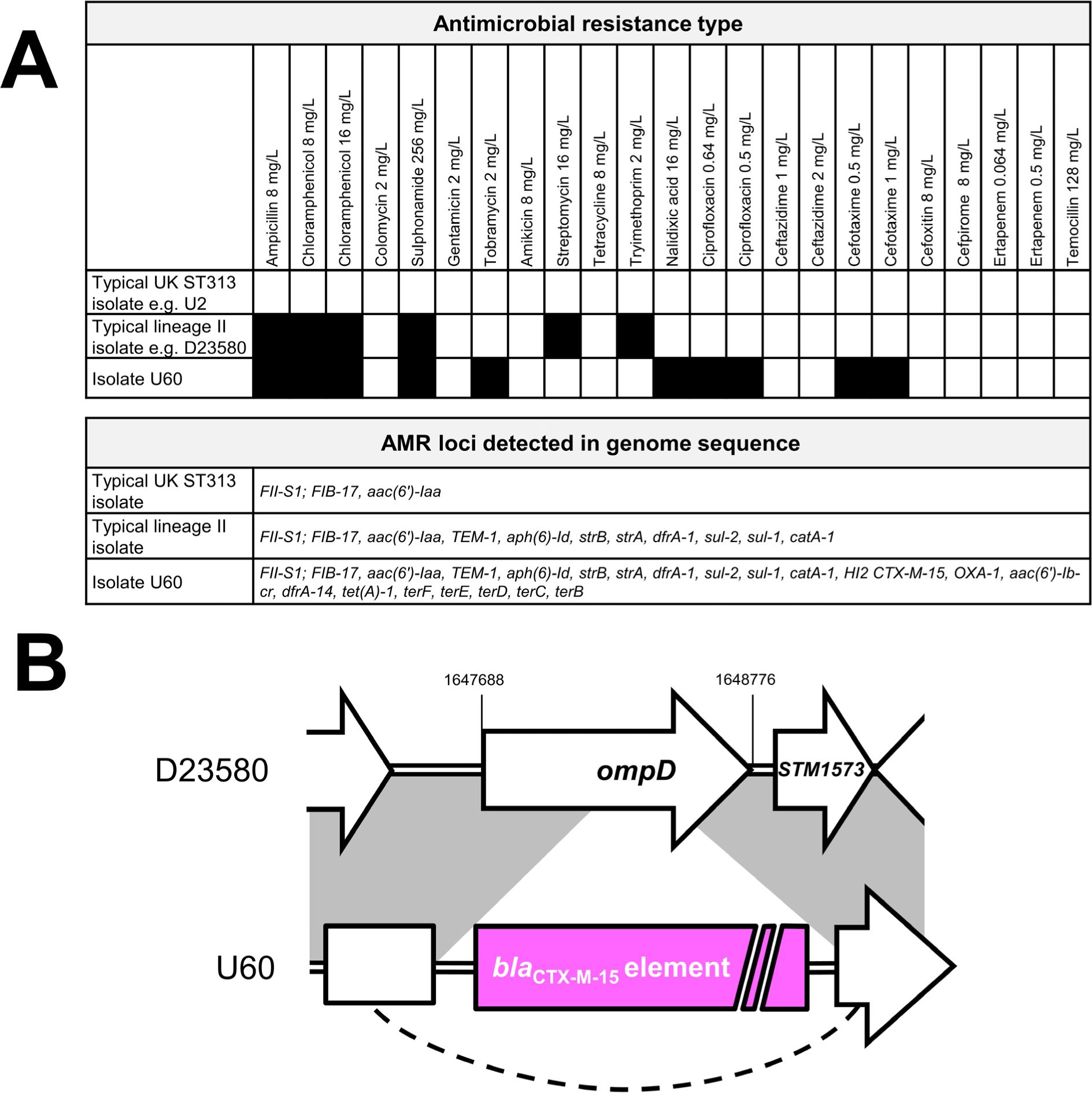
Isolate U60 contains additional resistance genes including a *bla*_CTX-M-15_ locus inserted into the chromosomal *ompD* locus. A. Antimicrobial resistance typing data and resistance genes detected in genome sequences for isolate U60, compared to data for typical lineage 2 and UK-ST313 isolates. B. Schematic illustrating the insertion of *bla*_CTX-M-15_ element into the chromosomal *ompD* locus in isolate U60. Further information is given in Supplementary Figure S1.

More detailed analysis of the genome of isolate U60 showed that a copy of the *bla*_CTX-M-15_ gene was inserted into the chromosome (location 1648104-1648109 on the D23580 reference genome), disrupting the *ompD* locus (Figure 4b Supplementary Figure S1). ESBL resistance genes have been reported previously in African ST313 isolates carrying plasmids such as pKST313 (21, 22) but this is the first report of a chromosomally-encoded ESBL resistance gene in *S.* Typhimurium ST313.

The assembled genomes of the UK-ST313 isolates were compared to the African ST313 reference strain D23580 using BLAST (Figure 2B). In agreement with published data (4), the majority of the core genome, including the *Salmonella* Pathogenicity Island (SPI) repertoire was conserved in the ST313 isolates in this study and in 3 ST19 gastroenteritis isolates (Figure 2B). The African ST313 lineages carry two prophages, BTP1 and BTP5, that are absent from ST19 strains (15). The entire BTP1 and BTP5 prophages were found in most ST313 isolates that belonged to African lineage 2 (12/13), but one UK-isolated lineage 2 strain, U68, lacked both prophages. The complete BTP1 and BTP5 prophages were not identified in any of the UK-ST313 isolates (Figure 2B), though some isolates contained partial and fragmented identity to BTP1 and BTP5, indicating the presence of related prophages (23, 24) which may not occupy the same attachment site. As expected, the *st313-td* gene (25) was carried by all twelve lineage 2 strains isolated from the UK that contained prophage BTP1. Only 1/51 UK-ST313 isolates contained the *st313-td* gene (isolate U76), where it was located on a prophage with 90% identity to BTP1.

To confirm the conservation of chromosomal organization in the UK ST313 isolates, representative isolate U2 was re-sequenced by PacBio long-read sequencing. The assembly produced two closed circular contigs representing the chromosome and the pSLT virulence plasmid (Supplementary Figure S2). Comparison with the ST313 lineage 2 reference genome D23580 identified no large chromosomal re-arrangements, deletions or duplications, and confirmed that the BTP1 and BTP5 attachment sites were unoccupied and did not contain additional prophages. No additional plasmids larger than the detection limit of the PacBio sequencing (~10kb) were detected in isolate U2.

### Genome degradation and pseudogenes in UK and African ST313

The ST313 lineages 1 and 2 responsible for iNTS disease in Africa have undergone genome degradation (6, 8). The pseudogenes identified in lineage 2 representative strain D23580 (6) were put into the context of the high-quality finished genome of UK-ST313 isolate U2 (Figure 5). The majority (34/44) of pseudogene mutations were conserved in U2. The only pseudogenes associated with characterized genes present in lineage 2 but functional in UK-ST313 strain U2 were *macB*, *ssel* and *lpxO* (Figure 5).

**Figure 5.**
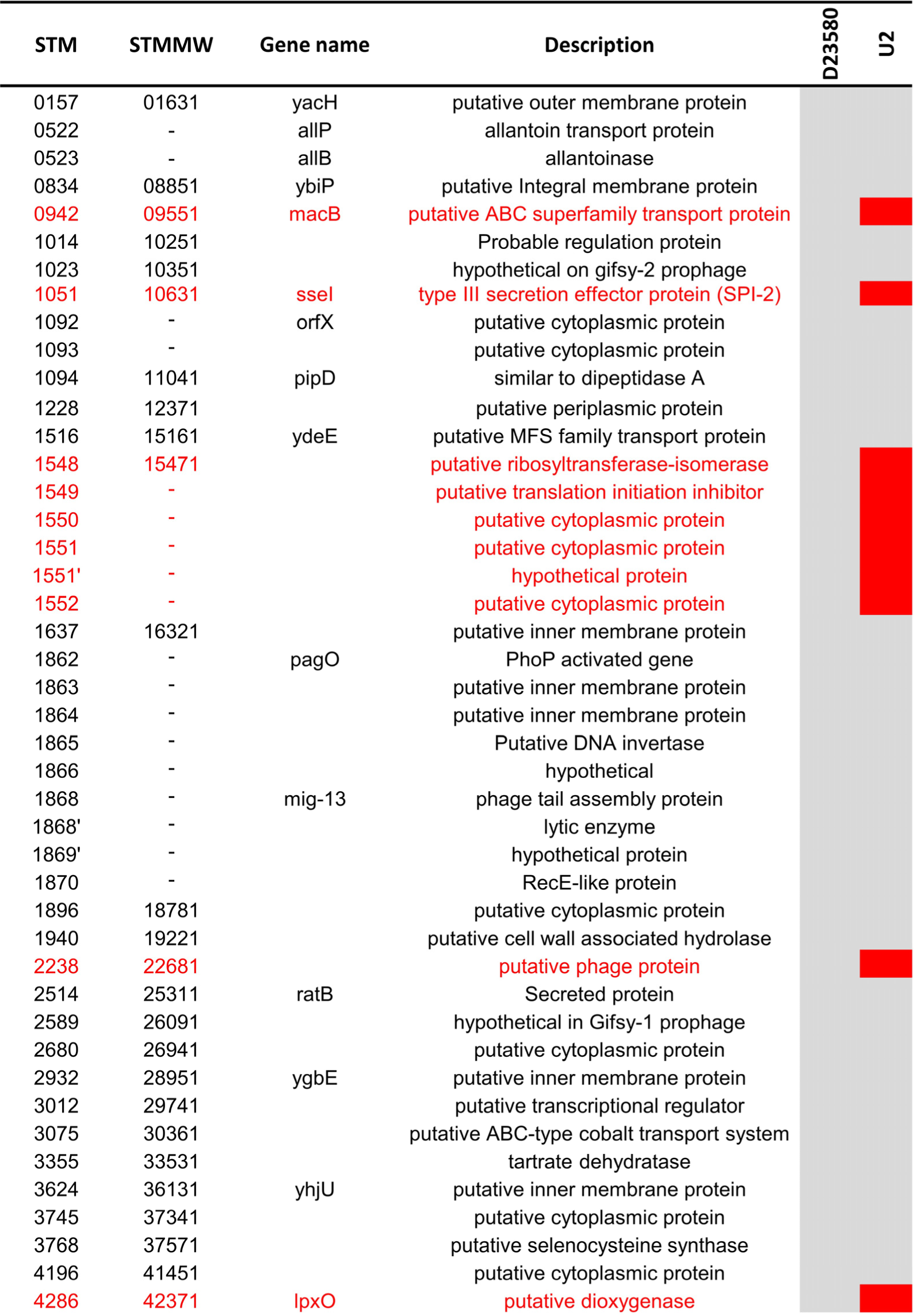
The majority of pseudogenes identified in lineage 2 strain D23580 are conserved in UK-ST313 representative strain U2. Heat map adapted from Kingsley *et al.*, 2009 showing genome degradation in ST313 strain D23580 (first heat map column) in the context of strain U2 (final column). Grey indicates pseudogenes conserved in both strains, whilst red indicates genes which are not degraded, and therefore likely functional, in strain U2.

### Phenotypic characterization of a subset of UK-ST313 strains

Several studies have associated the ability of ST313 to cause iNTS disease with particular phenotypic characteristics, such as the lack of RDAR morphotype formation, reduced swimming motility and the inability to produce catalase at stationary phase(18, 19, 26). We investigated these phenotypic characteristics in the context of the UK-ST313 strains, using a subset of 16 UK-isolated ST313, consisting of 13 UK-ST313 isolates and 3 lineage 2 isolates. The phylogenetic context of these 16 isolates is shown in Supplementary Figure S3. Lineage 2 representative strain D23580 and ST19 representative strain 4/74 were included as positive and negative controls.

The swimming motility of UK-isolated ST313 was highly variable between isolates (Figure 6a). One lineage 2 strain, U1, showed low levels of motility. However, this strain was observed to have a growth defect (data not shown). The ST313 lineage 2 representative strain D23580 was less motile than ST19 strain 4/74, consistent with previous reports (18). However, there was no apparent association between motility (as measured by migration diameter) and phylogenetic context of the lineage 2 strains and the UK-ST313 strains.

**Figure 6.**
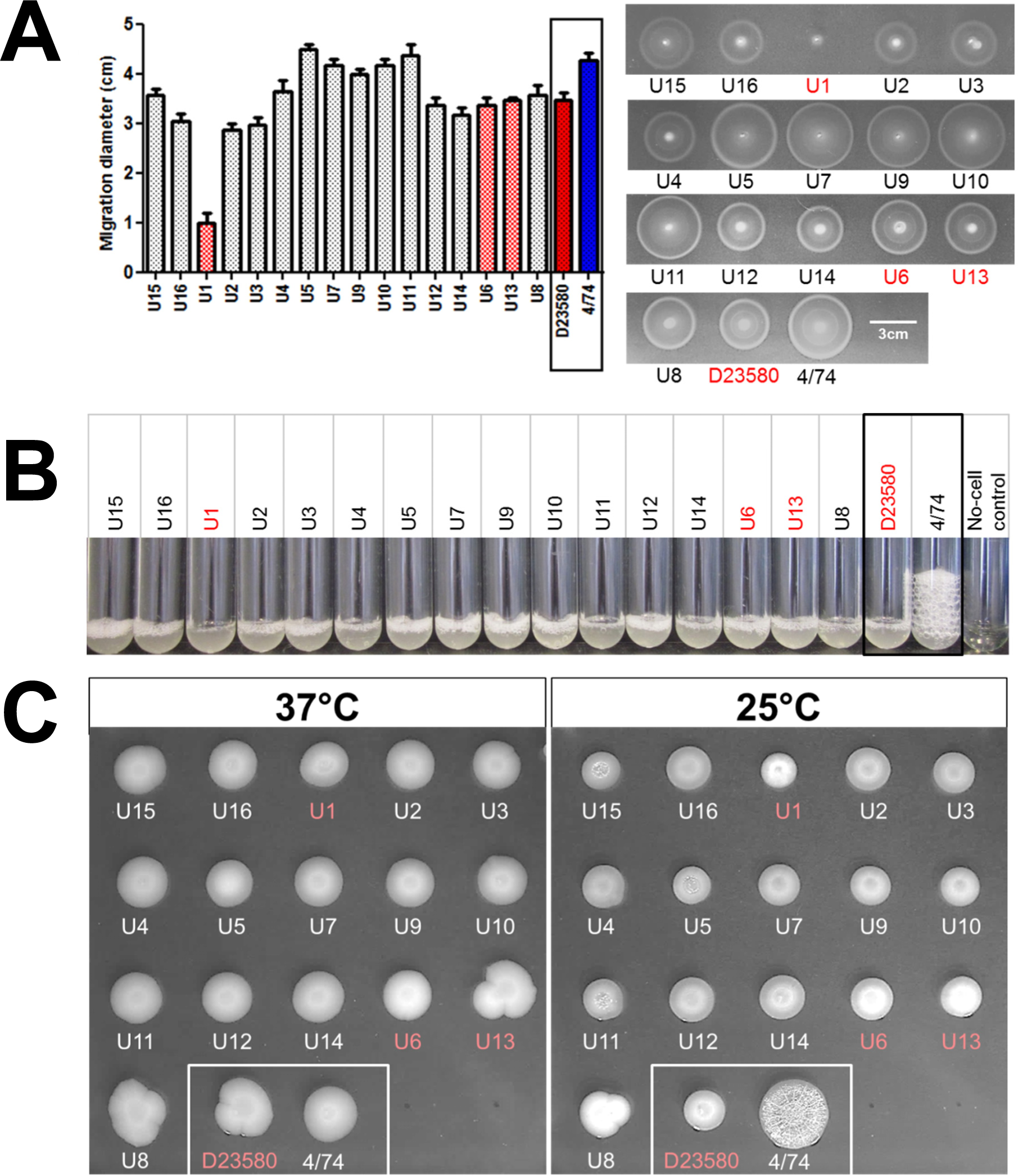
*In vitro* phenotypes of a subset of UK-isolated ST313 strains in the context of representative ST313 lineage 2 and ST19 strains D23580 and 4/74. UK-isolated strains that belong to African lineage 2 (U1, U6 and U13) are highlighted in red throughout. A. Migration diameter after 5 hours (average of 3 replicates is shown together with error bars representing standard deviation). A representative plate is shown, right. B. Stationary phase catalase activity represented by bubble column height after 5 minutes exposure to 20 µl 20% H_2_O_2_. C. RDAR morphology assay. RDAR phenotype forms after prolonged incubation at 25°C but not at 37°C.

The *katE* pseudogene was reported to be responsible for the lack of catalase activity in ST313 lineage 2 (19). All 16 UK-isolated strains were shown to be negative for stationary phase catalase activity, as was the lineage 2 representative strain D23580 (Figure 6b). In contrast, ST19 strains 4/74 showed considerable stationary phase catalase activity, consistent with previous findings (19).

The RDAR morphotype of *Salmonella enterica* is linked to resistance to desiccation and exogenous stresses (27). African lineage 2 ST313 are reported to be defective in RDAR morphotype formation due to a truncated BcsG protein due to the introduction of a premature stop codon (19). All the UK-isolated strains and the African lineage 2 reference strain D23580 did not exhibit the RDAR morphotype. In our experiments, only the ST19 strain 4/74 exhibited the RDAR morphotype (Figure 6c).

These experiments did not identify any phenotypic differences between the UK-ST313 isolates and ST313 lineage 2, and future work is needed to identify African ST313-specific phenotypes.

## Discussion

Recent reports of iNTS disease have been associated with the novel *Salmonella* Typhimurium ST313 in sSA (6, 7, 16), and suggested that ST313 was geographically restricted to sSA. This prompted the investigation of the presence of this sequence type amongst *S*. Typhimurium isolates from the UK.

We discovered that 2.7% of *Salmonella* Typhimurium isolates referred to Public Health England are of MLST type ST313, and that this sequence type is heterogeneous in terms of clinical presentation, genomic characteristics and epidemiology. The UK-isolated ST313 strains are predominantly fully antimicrobial-susceptible and cause gastroenteritis. We identified a significant association between travel to Africa and infection with the previously described African-associated, ST313 lineage 2. We found two cases of lineage 2 infection where the patient did not report travel to Africa, although one of these patients reported consumption of food from West Africa in London. This indicates that African lineage 2 is predominantly circulating in Africa but may also be circulating in the UK and other countries – potentially via person to person transmission or through exposure to food imported from Africa. People infected with ST313 lineage 2 in the UK were significantly more likely to suffer from invasive disease than patients infected with isolates belonging to UK-ST313 lineages. There is also an increasing number of immune-compromised people in the UK with increasing organ transplants and immune-modulating cancer therapies.

This study revealed novel diversity within ST313, which was previously restricted to two African lineages that had exhibited recent clonal expansion (7). Here we place these lineages into an evolutionary context by showing that lineage 1 and 2 do not form a monophyletic group within ST313, which is suggestive of two separate introductions of ST313 into sSA. African lineages 1 and 2 diverged from their MRCA with UK-lineages around 1796 and 1903 respectively. These findings reflect the limitations of classifying bacterial pathogens simply on the basis of sequence type, and show that in the post-genomic era, the resolution offered by MLST may not be sufficient to describe epidemiologically relevant population structures.

It has been estimated that 9.2% of cases of Salmonellosis in the EU can be attributed to international travel, and therefore sequencing *Salmonella* isolated in Europe can provide valuable information regarding the global diversity of *Salmonellae* associated with human disease (28, 29). The genome of one UK-isolated lineage 2 isolate associated with travel to Kenya, U60, contained sequences with high nucleotide similarity to a recently described IncHI2 plasmid, pKST313, associated with Kenyan ceftriaxone resistant ST313 isolates (21). The U60 plasmid encoded 4 additional MDR genes and tellurium heavy metal resistance genes. Until now the *bla*_CTX-M-15_ gene has only been found to be plasmid-associated in Salmonellae. We discovered that the *bla*_CTX-M-15_ gene was chromosomally encoded in isolate U60, causing disruption of the *ompD* locus. This is notable for two reasons. Firstly, chromosomal integration ensures stability of ESBL-resistance even if the plasmid were lost. Secondly, *ompD* encodes an outer membrane porin that is absent from *Salmonella* Typhi. OmpD has previously been identified as a highly immunogenic protein (30) and so the disruption of *ompD* could enhance the reported “stealth” phenotype of ST313 lineage 2 infection (17). We note that OmpD is a potential vaccine target for iNTS (31) and the absence of OmpD from African ST313 populations could have implications for future iNTS vaccine development.

Our discovery of UK-ST313 isolates that were not associated with invasive disease provides an excellent opportunity to use comparative genomics to relate genetic findings that have been linked the pathology of lineage 2 ST313 into the context of closely related, gastrointestinal-associated strains. We found that the only genetic characteristics common to both lineages 1 and 2 and absent from the UK-ST313 genomes were the BTP1 and BTP5 prophages and plasmid-associated MDR loci. The two African lineages do not share a common ancestor that carried either prophage, suggesting independent acquisition of BTP1 and BTP5 by lineage 1 and 2, and whilst the MDR loci confer similar patterns of AMR they are genetically distinct. This implies a strong selection pressure on ST313 in Africa to acquire and maintain these mobile elements, resulting in convergent evolution of the two African lineages. In contrast, there is evidence of an assortment of distinct prophage repertoires in the UK-ST313 isolates, indicating an absence of selection for the conservation of any particular mobile element.

Aside from the addition of mobile genetic elements and virulence factors, genome degradation by the accumulation of pseudogenes and deletion events is known to accompany adaption to a more invasive lifestyle (32, 33). Initial analysis of the African ST313 representative strain D23580 genome reported 23 pseudogenes compared to 6 present in ST19 strain SL1344 (34). Here, we found that the majority of genome degradation found in lineage 2 strain D23580 was conserved in UK representative strain U2. The only pseudogenes associated with characterized genes that were found to be specific to African lineage 2 ST313 were the SPI2-secreted effector gene *ssel*, lipid A modification gene *lpxO* and macrolide efflux pump gene *macB*, each of which could play a role in infection dynamics (35–37).

A number of *in vitro* phenotypes have been reported for lineage 2 ST313, that could contribute to a host-adapted lifestyle (17–19, 26) and were examined in the UK-ST313. Swimming motility was highly variable amongst the strains tested, and UK-ST313 isolates behaved identically to African lineage 2 isolates in the catalase and RDAR morphotype assays. We detected no African-lineage-specific phenotypic characteristics and speculate that reduced motility, defective catalase activity and loss of RDAR formation are not be directly linked to iNTS disease.

A key contributing factor to iNTS disease is host immunosuppression and one limitation of this retrospective study was that the underlying health status of the patients was unknown. This study does highlight the extraordinary epidemiological insights that routine genomic surveillance of pathogens by public health agencies can offer, and the ability to understand the pathogenesis of novel pathovars. The knowledge that an immune-compromised patient was infected with lineage 2 ST313 could impact clinical decision-making.

We have uncovered previously un-sampled diversity in the ST313 clone reflecting the convergent evolution towards niche specialization that has occurred in the African lineages. The routine genomic surveillance of pathogens continues to be adopted internationally and will bring an unprecedented ability to monitor emerging threats. Whole genome sequencing of clinical isolates represents a new window to view the epidemiology and microbiology of infectious diseases.

## Materials and Methods

### Strains and Metadata

Genome sequences from a total of 363 *Salmonella* Typhimurium isolates dating from 2012 and 3,014 *Salmonella* Typhimurium from January 1^st^ 2014 to March 14th 2016 were analyzed for this study.

For simplicity, isolates derived from blood culture, the infection was classed as extra-intestinal. If only a faecal isolate was received for a patient, the infection was classed as gastrointestinal (though this is only suggestive, not conclusive, data that the infection was restricted to the gastrointestinal tract). Of these, 7/363 (1.9%) and 79/3,014 (2.6%) were *Salmonella* Typhimurium ST313, respectively. Full strain metadata can be found in Table S1. Sequence data (FASTQs) from 23 representative ST313 sequenced by Okoro *et al.* (7) were downloaded from the European Nucleotide Archive (accessions available in Table S1) and analyzed in the same way as sequence data generated from UK-isolated ST313 isolates.

### Sequencing

DNA extraction for Illumina sequencing of *Salmonella* isolates was carried out using a modified protocol of the Qiasymphony DSP DNA midi kit (Qiagen). In brief, 0.7 ml of an overnight *Salmonella* culture in a 96 deep well plate was harvested. Bacterial cells were pre-lysed in 220 μl of ATL buffer (Qiagen) and 20 μl Proteinase K (Qiagen), and incubated with shaking for 30 mins at 56°**C**. Four microliters of RNase (100 mg/ml; Qiagen) was added to the lysed cells and re-incubated for a further 15 minutes at 37°C. This step increased the purity of the DNA for downstream sequencing. DNA from the treated cells was then extracted on the Qiasymphony SP platform (Qiagen) and eluted in 100 μl of sterile water. DNA concentration was derived using the GloMax system (Promega) and quality (optimal OD260/230 = 1.8 − 2.0) was determined using the LabChip DX system (Perkin Elmer). Extracted DNA was prepared using the NexteraXT sample preparation method, and sequenced with a standard 2×100 base pair protocol on a HiSeq 2500 instrument (Illumina, San Diego). Raw FASTQs were processed with Trimmomatic (38) and bases with a PHRED score of less than 30 removed from the trailing end.

PacBio sequencing was performed on the PacBio RS II instrument at the Centre for Genomic Research, University of Liverpool. DNA was extracted from strain U2 using the Zymo Research Quick-DNA^TM^ Universal Kit (cat# D4069) as per the Biological Fluids & Cells protocol. Extracted DNA was purified with Ampure beads (Agencourt) and the quantity and quality was assessed by Nanodrop and Qubit assays. In addition, the Fragment analyzer (VH Bio), was used to determine the average size of the DNA, using a high sensitivity genomic kit. DNA was sheared to approximately 10kb using a Covaris g-tube and spinning at 5400rpm in an Eppendorf centrifuge. The size range was checked on the Fragment Analyzer. DNA was treated with exonuclease V11 at 37°C for 15 minutes. The ends of the DNA were repaired as described by Pacific Biosciences. Samples were incubated for 20 minutes at 37°C with damage repair mix supplied in the SMRTbell library kit (Pacific Biosciences). This was followed by a 5-minute incubation at 25°C with end-repair mix. DNA was cleaned using 1:1 volume ratio of Ampure beads and 70% ethanol washes. DNA was ligated to adapters overnight at 25°C. Ligation was terminated by incubation at 65°C for 10 minutes followed by exonuclease treatment for 1 hour at 37°C. The SMRTbell library was purified with 1:1 volume ratio of Ampure beads. The library was size-selected on the Blue Pippin (Sage) in the range 7kb-20kb. The DNA was recovered and the quantity of library and therefore the recovery was determined by Qubit assay and the average fragment size determined by Fragment Analyser. SMRTbell libraries were annealed to sequencing primer at values predetermined by the Binding Calculator (Pacific Biosciences) and complexes made with the DNA polymerase (P6/C4 chemistry). The complexes were bound to Magbeads and loaded onto 3 SMRT cells. Sequencing was done using 360-minute movie times. Sequence data from the 3 SMRT cells was assembled using the HGAP3/Quiver assembler. This resulted in 2 contigs representing the chromosome and the pSLT virulence plasmid. Terminal repeats were manually trimmed to represent circular molecules, and the chromosome assembly was reordered so that the sequence started at the *thrL* locus in accordance with convention for *Salmonella* finished genomes. The closed sequences for the U2 chromosome and pSLT virulence plasmid were 4,811,399bp and 93,862bp respectively. Prokka (39) was used to annotate the two sequences, using the –force flag to preferentially annotate CDS from reference databases FN424405 for the chromosome and AE006471 for the virulence plasmid. The finished U2 genome and annotation were submitted to Genbank and can be accessed using the study number PRJEB20926.

### Genomic analysis

The multi-locus sequence type (ST) was determined using a modified version of SRST (40). For phylogenetic analysis, processed sequence reads were mapped to the *S*. Typhimurium LT2 reference genome (GenBank: AE006468) using BWA mem (41). SNPs were called using GATK2 (42) in unified genotyper mode. Core genome positions that had a high quality SNP (>90% consensus, minimum depth 10x, MQ >=30) in at least one strain were extracted and IQ-TREE with parameters –m TEST – bb 1000 was used to construct a maximum likelihood phylogeny (43).

To examine the evolutionary history of ST313, four timed phylogenies were constructed using BEAST v1.8.0 (44), with varying clock rate models and tree priors. The resulting models were compared in terms of their tree likelihood and posterior and the strict exponential and strict constant models were found to be superior. A comparison using AICM calculated with Tracer v1.6.0 showed that the models had very similar values, tree topologies and branching support in terms of posterior probability were similar between the models. The 95% HPD for the exponential growth rate estimate was −0.0026 to 0.0006; the strict, constant growth model was selected as the estimate of growth rate from the exponential model was around 0 (i.e. constant).

Accessory genome analysis was performed using *de novo* assemblies of quality processed FASTQs produced using SPAdes v2.5.1 using default parameters except –careful and –k 22, 33, 55, 77 (45). Searching for specific gene targets e.g. *st313-td* was performed using BLAST+ within the BioPython framework (46) and whole genome assemblies were compared to the reference ST313 strain D23580 using BRIG (47).

### Microbiology

Phenotypic antimicrobial susceptibility testing was carried out for all UK-isolated ST313 strains. The antimicrobial susceptibility testing was done using breakpoint concentrations. Briefly, an agar dilution method involving Iso-sensitest agar or Muller-Hinton agar was used to determine if isolates were sensitive or resistant to a set concentration of individual antimicrobials (Supplementary Table S2).

Swimming motility assays were performed based on methods previously described (48). A 3µl aliquot of bacteria grown overnight in LB (Lennox Broth; 10 g/L Bacto Tryptone, 5 g/L yeast extract and 5 g/L NaCl, pH7.0) was spotted onto LB (Lennox) plates containing 0.3% Bacto Agar (Difco). Plates were incubated at 37°C. After exactly 5 hours the migration diameter was measured and plates were photographed.

Catalase activity and RDAR morphotypes were assayed based on methods used by Singletary *et al.* (19). Briefly, for catalase activity, 20 µl of 20% aqueous H_2_O_2_ was added to 1ml of bacteria grown overnight in LB (Lennox), in 1cm diameter glass test tubes. Tubes were photographed after 5 minutes incubation at room temperature and the height of the bubble column measured. For RDAR morphology, 2 µl of bacteria grown overnight in LB (Lennox) were spotted onto LB plates without NaCl and supplemented with 40 µg/ml Congo red and 20 µg/ml Coomassie blue. Plates were incubated at 25°C and 37°C for 7 days without inversion. All experiments were conducted in triplicate.

### Epidemiology

Food poisoning is a notifiable disease in the UK and diagnostic laboratories are obliged to report the isolation of *Salmonella* from human clinical diagnostic samples. However, data are frequently incomplete and detailed exposure information for cases is not always available in retrospect. Therefore, targeted surveillance questionnaires were attempted to obtain enhanced information, focusing primarily on collection of information on clinical severity of disease, travel history and consumption of foods of African origin were utilized during telephone interviews for cases reported from 2014-2016 to collate relevant epidemiological data. Cases for which enhanced information were available are shown in Table S1.

Collection of this epidemiological data was not attempted for the 2012 cases, but limited travel data had been recorded on the SRS *Salmonella* surveillance database or some isolates. It is important to emphasize that the travel information for the 2012 isolates is of low quality, and the absence of reported travel does not mean that international travel had not occurred. Odds ratios were calculated using the medcalc.org website https://www.medcalc.org/calc/odds_ratio.php.

## Acknowledgements

We would like to acknowledge PHE Genomic Services and Development Unit, Infectious Disease Informatics and *Salmonella* Reference Service members, in particular Martin Day for antibiotic resistance typing. We are grateful to John Kenny, Margaret Hughes and Xuan Liu at the Centre for Genomic Research, University of Liverpool for assistance with PacBio sequencing. We would also like to thank Rob Kingsley and Rocío Canals for helpful discussion and comments.

